# Physical confinement and phagocytic uptake induce directional migration

**DOI:** 10.1101/2025.05.13.653701

**Authors:** Summer G. Paulson, Sophia Liu, Jeremy D. Rotty

## Abstract

Physical confinement is not routinely considered as a factor that influences phagocytosis, which is typically assayed in unconfined settings *in vitro*. BV-2 microglia-like cells were used to interrogate the impact of confinement on IgG-mediated phagocytosis side by side with unconfined cells. Confinement acted as a potent phagocytic driver, greatly increasing the fraction of phagocytic cells in the population compared to the unconfined setting. Arp2/3 complex and myosin II contributed to this effect. Remarkably, confinement partially rescued phagocytic uptake upon myosin II disruption. In addition, cells under confinement were partially resistant to the actin-depolymerizing drug cytochalasin D. Unexpectedly, we observed that bead uptake stimulated persistent migration, a process we term ‘phagocytic priming’. Integrin-dependent adhesion was required for phagocytic priming in unconfined and confined settings, but was dispensable for phagocytic uptake. The cytoskeletal requirements for phagocytic priming differed depending on confinement state. Myosin II and Arp2/3 complex were required for phagocytic priming under confinement, but not in unconfined settings. As with phagocytosis, cytoskeleton-dependent priming of motility shifts depending on physical confinement status. Phagocytic priming may facilitate innate immune function by driving cells to more efficiently surveil their local microenvironment in response to wounds or trauma.

## INTRODUCTION

Many microenvironmental factors act as external stimuli to influence cell morphology or function (*1*, *2*). These can be chemical cues that elicit directed cell migration, like chemotactic or haptotactic gradients, or physical cues, such as tissue stiffness and physical confinement that alter cell shape or modify cellular responses. Confinement is defined as any external cue that restricts cell morphology or mobility (*3*, *4*). These microenvironmental factors and others combine to influence a cell’s response to its environment and with other cells around it, such as prompting a phagocytic response or shifting motility (*1*, *2*).

The physical microenvironment is sufficient to alter cellular responses. For example, cells reintroduced into decellularized extracellular matrix (ECM) structures assume identities similar to the missing cell population (*5*). Hydrogels derived from decellularized ventricular ECM increased the number of endogenous cardiomyocytes in rats (*6*) and decellularized, demineralized bone matrices bonded with vascular endothelial growth factor increased endothelial cell proliferation and promoted angiogenesis via new microvessel invasion within the scaffolds (*7*). A study on the transfer of decellularized ECM to new surfaces, facilitating expansion and maintenance of mesenchymal stem cells, demonstrated that the component used in the ECM transfer process (urea, pepsin, or acetic acid) had an effect on how the stem cells responded to the new ECM (*8*).

Microglia, the resident immune cell of the central nervous system (CNS), are responsible for maintaining CNS homeostasis by supporting synapse formation, clearing debris, and responding to inflammatory cues within the brain and spinal cord (*9–12*). The brain itself is a highly confined space (**Figure 1A**) (*13*), providing an example for the need to factor microenvironmental cues into *in vitro* assays when emulating specific tissues. However, it is difficult to examine the effects of confinement and other microenvironmental factors on cellular behavior in the brain, where decellularizing and reinjecting ECM matrix is less feasible. In addition, microglia have a defined ramified phenotypic morphology *in vivo,* and microglia *in vitro* often do not maintain that established morphology, trending instead towards more amoeboid-like morphologies (*14*, *15*). Amoeboid cells are more migratory and active than ramified ones, and therefore more likely to phagocytose (*16*). This points towards *in vitro* assays lacking microenvironmental cues that replicate the ramified morphology and activity of microglia *in vivo*. A recent study by Sharaf *et al.* utilized nano-pillar arrays as a 2.5D assay for microglial growth and recovered the ramified cell morphology compared to amoeboid-like cells in standard 2D culture (*17*). However, this study used relatively hard surfaces for their pillars with a Young’s modulus of 0.25-11.4MPa. The Young’s modulus of brain tissue is significantly softer, ranging from 0.1-1kPa (*18*, *19*). A common confinement strategy utilized in chemotaxis studies confines cells under 1% agarose, which has a stiffness in the same range as brain (*20*, *21*).

**Figure 1:**
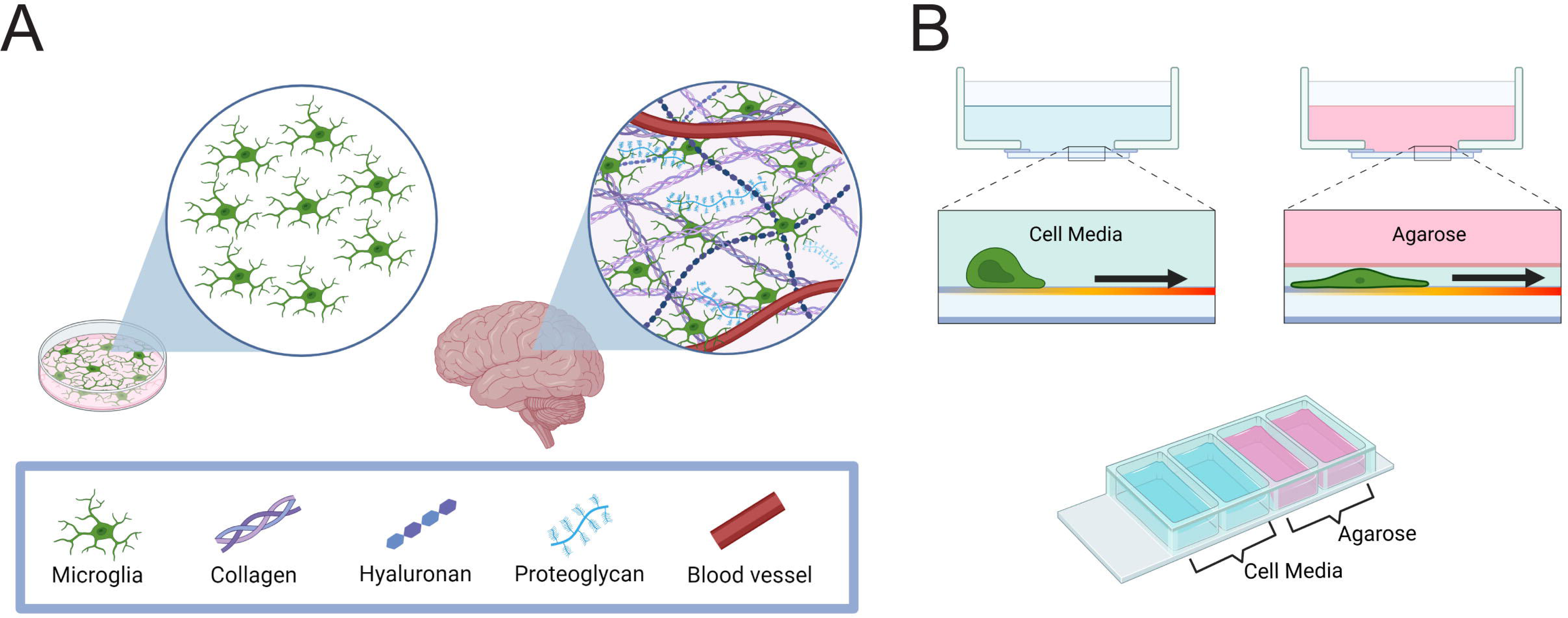
The brain is a confined environment. A) Schematic demonstrating microenvironmental components confining cells in the brain, compared to cells in a petri dish that are allowed to freely move B) Top: Examples of cells moving with and without a confining layer of agarose. Bottom: schematic of cell assay used throughout. A 4-chamber well glass bottom dish was coated with 10µg/mL fibronectin and then filled with either complete cell media or 1% agarose. Cells were introduced into wells and allowed to move.

Elastic, soft confining environments composed of agarose, collagen, or Matrigel have been used to interrogate the role of the actin cytoskeleton in cellular behavior. A study done with dendritic cells under agarose demonstrated that increased agarose stiffness resulted in a decrease in cell velocity and distance travelled during cell migration (*22*). With heightened agarose stiffness, the cells created actin patches, utilizing WASp and the Arp2/3 complex to facilitate pushing the actin patches against the plasma membrane to raise the confining environment upward so that the cell could propel itself forward. A study with cells plated on polyacrylamide gels demonstrated that the introduction of fibronectin micropatterns resulted in increased adhesion and prevented cell spreading, instead confining the cells to the fibronectin micropatterns (*23*). They found that cells in these fibronectin patches on the polyacrylamide gel had an altered flow of F-actin within the cells, moving in a retrograde fashion instead of outward to contain the cells to the adhesion patch. Confining macrophages with either a micropattern or by cell crowding downregulated the pro-inflammatory cytokine secretion that precedes phagocytosis (*24*). Another study done with bone marrow derived macrophages in Matrigel found that inhibition of F-actin, ROCK, or non-muscle myosin II (NMII) decreased cell migration speed compared to untreated confined cells (*25*). Each of these results pointed towards the importance of several components of the actin cytoskeleton in confined migration. But, limited research has been done reading how phagocytosis occurs in a confined environment.

Therefore, we set out to examine the effects of confinement on microglia phagocytosis, utilizing agarose with a stiffness akin to brain parenchyma, by directly comparing IgG phagocytosis in confined and unconfined cells. We discovered that confinement promotes more efficient phagocytosis, which is dependent upon Arp2/3 complex and non-muscle myosin II. We also noted that confinement partially rescues the effect of myosin II inhibition, as well as cytochalasin D treatment. These findings imply that there may be an actomyosin-independent phagocytic component that becomes activated under physical confinement. We also discovered that confinement and phagocytic uptake work together to induce what we define here as ‘phagocytic priming’, in which cells migrate in a highly persistent fashion immediately after phagocytosis.

Phagocytic priming required integrin-dependent adhesion to the ECM in confined and unconfined settings, while Arp2/3 complex and myosin II function were only required under confinement. As a whole, this work supports the notion that the molecular requirements for phagocytosis and cytoskeleton-dependent priming may be context-dependent, and may be especially responsive to physical confinement. In addition, phagocytic priming may be crucial for supporting innate immune processes like homing of antigen-presenting cells to the lymph node, for enhanced exploration of the microenvironment, or for phagocytic responses elicited by wounds or infection.

## RESULTS

### Confinement enhances phagocytic uptake and persistent migration

Much of what has been established regarding the molecular mechanisms of glial cell biology has been established in a 2D culture system. Therefore, an assay was designed to be able to compare the influence of confinement on cells alongside their response to phagocytic ligands in a confined setting. Using a four-chamber dish, cells could either be placed in a well filled with media or confined under a layer of 1% agarose (**Figure 1B, bottom**). These assays are based on previous cell motility experiments in the lab (*26*), in turn inspired by the classic under-agarose chemotaxis assay as described by Heit and Kubes (*20*). This created the confinement-induced restriction on cell function as they move in response to a cue (**Figure 1B, top**).

First, we compared the ability of microglia like BV-2 cells to phagocytose either in media or confined under 1% agarose. Murine IgG was labeled with a pHrodo fluorescent tag and used to opsonize 2µm polysphere beads. The pHrodo tag was used because it fluoresces under acidic environments, like the phagolysosome. Thus, we were able to detect complete phagocytosis in an unbiased fashion over 8-hour time lapse movies (**Supplemental Figure S1A**). To rule out any signal from external beads, we compared external and internalized beads and demonstrated that the difference between these two states is clear (**Supplemental Figure S1B)**. In order to normalize the number of beads present when comparing between media and confined wells, bead densities were measured and categorized into different density groups (**Supplemental Figure S1C-D**). Beads injected under agarose did not disperse as evenly as in media wells. Due to this variability between fields of view under confinement, the number of beads were calculated to be able to control for bead density when comparing confined and media wells. Based on the majority of media well fields of view being categorized as ‘low bead density’ (**Supplemental Figure S1D, bottom**), fields of view within this matched ‘low bead density’ under confinement (**Supplemental Figure S1D, middle)** were exclusively used for analysis.

Phagocytosis was measured at the 2-hour mark, before cells became saturated with beads and stopped phagocytosing. Example images for both the media well (**Figure 2A**) and the confined well (**Figure 2B**) demonstrate pHrodo fluorescence signal from internalized beads. Cells under confinement were significantly more phagocytic than unconfined (media) cells (**Figure 2C**). While more cells became phagocytic under confinement, there was no overall difference in phagosome size or number when comparing pHrodo^+^ cells in either condition (**Figure 2D-F**). As a whole, this data demonstrates that confinement induces a higher number of cells to phagocytize but does not affect the efficiency of intracellular trafficking to the phagosome after bead uptake.

**Figure 2:**
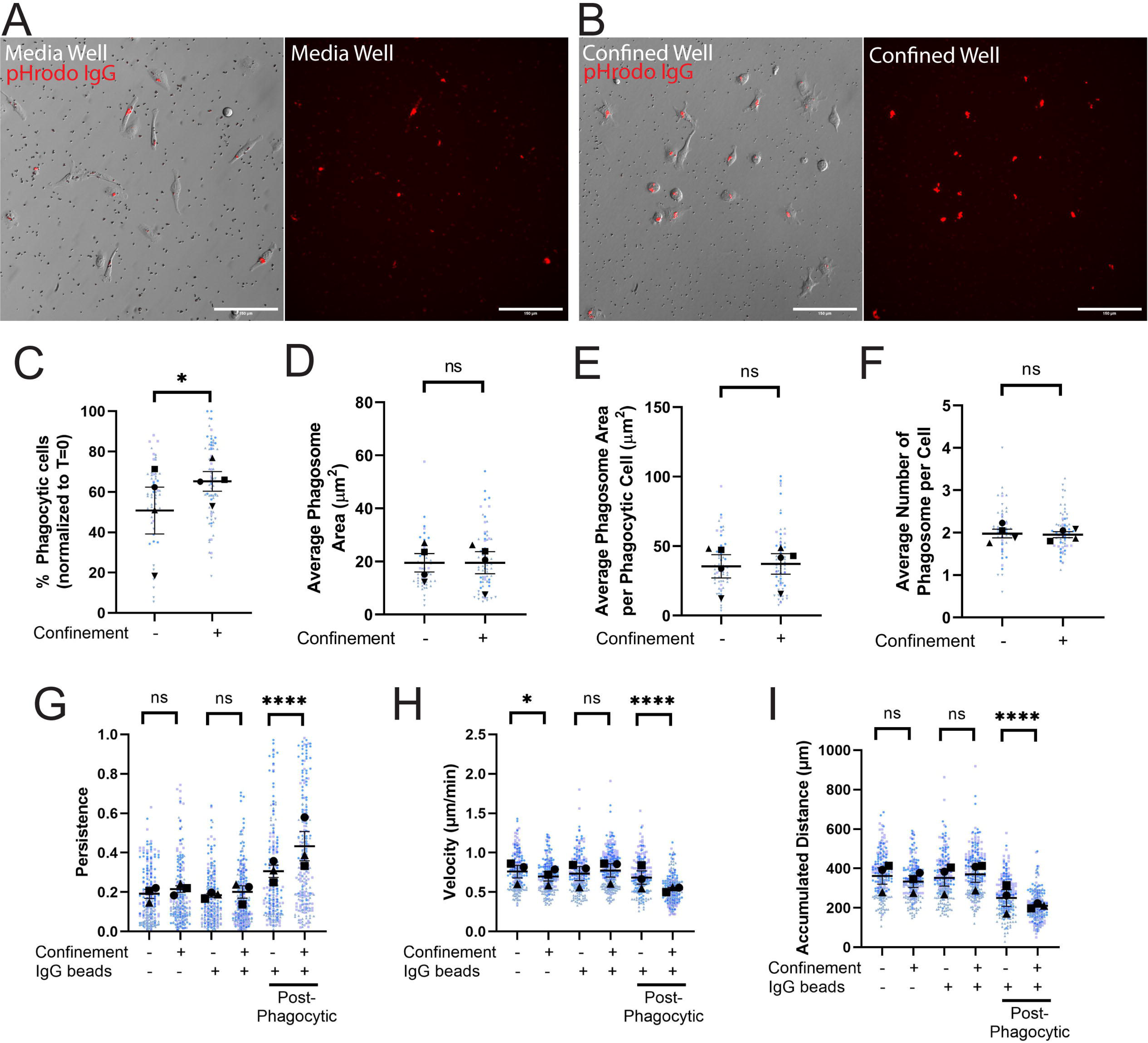
Confinement enhances phagocytosis and primes post-phagocytosis cells to migrate persistently. A-B) Example composite images of phase contrast and pHrodo-red merged images 2 hours after pHrodo-IgG beads were added to media wells (A) or confined agarose wells (B). The pHrodo-red signal from these merges has been isolated alongside the merge to emphasize internalized pHrodo-IgG-bead signal. Scale bar represents 100µm. C) The percentage of fluorescent cells in a field of view, comparing unconfined and confined cells, normalized to T=0. Any cell fluorescent at T=0 was removed from further counting. D) Average phagosome size (µm^2^). This is the average size of phagosomes per field of view divided by the average number of phagosomes per field of view. E) Average phagosome area per phagocytic cell (µm^2^). This was calculated by dividing the average size of all phagosomes in a field a view by the number of fluorescent cells counted (i.e. phagocytic cells). F) Average number of phagosomes per phagocytic cell. G) The persistence of the cell during the length of its track. X-axis labels denote whether the cells were in media or confined and if there were beads present in the well. The last two columns were the same fields of view as counted in the middle two columns, but tracking was conducted only after a cell phagocytized at least one pHrodo-IgG-bead. This also applies for H and I. H) Velocity of the cells migrating (µm/min). I) The maximum accumulated distance (µm) that the cell traveled over the track. For all graphs, black points demonstrate experiment means, colored points demonstrate individual cell values for each run. N = 4 experiments for each graph; n = 15 fields of view for each condition per experiment for phagocytosis (C-F) or n = 50 cells for each condition per experiment for migration (G-I). For (C-F), statistical analysis was assessed using the Mann–Whitney tests: ns = not significant, *p < 0.05. For (G-I), statistical analysis was assessed using via Kruskal–Wallis test with Dunn multiple comparisons: ns = not significant, *p < 0.05, ****p < 0.0001. Error Bars represent SEM.

While watching the time lapse movies, it became apparent that phagocytosis altered the migratory trajectory of cells. Specifically, bead uptake seemed to induce more persistent migration tracks compared to non-phagocytic cells (**Supplemental Movie 1**). Therefore, we decided to investigate whether bead uptake primed cells to move directionally. We measured cell velocity, accumulated migration distance, and migratory persistence. Persistence (D/t) is a measurement of a cell’s track from start to finish (t) and its net displacement during that timeframe (D), with values closer to 1 indicating more directionally persistent migration.

Increasing confinement pressure by placing cells under higher-percentage agarose gels led to decreasing velocity and accumulated distance, but no coherent persistence trend (**Supplemental Figure S2A-C**). With these data in hand, we returned to the setup from **Figure 1C**, except that each condition (media and confined) had one well with cells alone and one well with cells plus beads. We tracked cells across 8 hours and compared the cell velocity, accumulated distance, and cell persistence between groups. Notably, the -IgG beads condition revealed only a confinement-induced decrease in migration velocity (**Figure 2G-I**). When considering the +IgG beads as a bulk population (i.e. not separating out post-phagocytosis cells), we likewise saw no significant confinement-induced migration trends (**Figure 2G-I**). However, substantial differences were revealed when we narrowed our analysis only to the post-phagocytic phase. Confined and unconfined post-phagocytic cells migrated slowly, but with higher persistence than non-phagocytic cells (compare -IgG populations to post-phagocytic populations, **Figure 2G-I**). Furthermore, confined post-phagocytic cells were significantly more persistent, but slower, than even unconfined post-phagocytic cells (**Figure 2G-I**). This data points towards a process we have termed ‘phagocytic priming’, where phagocytosis stimulates directionally persistent migration. Confinement further increases persistent migration, enhancing the phagocytic priming response. Our next goal was to determine the cytoskeletal regulators involved in phagocytic priming and phagocytosis under confinement.

### The Arp2/3 complex is required for phagocytic priming and uptake

We first interrogated the Arp2/3 complex, a seven-subunit branched actin nucleator responsible for generating the branched actin networks that form lamellipodial protrusions linked to environmental sensing (*27*). While the Arp2/3 complex has been implicated in phagocytosis in unconfined settings (*28*), there is less known about its function during confined phagocytosis. Since the Arp2/3 complex is responsible for creating lamellipodial structures at the leading edge of polarized cells (*29*), we hypothesized that the Arp2/3 complex would be integral to both phagocytic uptake and priming under confinement.

We reutilized our 4-chambered dish, and this time added the Arp2/3 complex small molecule inhibitor CK-666 or vehicle control (DMSO) into the media or the agarose gel (**Supplementary Figure S2D**). Trial runs were done comparing cells with and without DMSO treatment to determine whether treatment with vehicle altered normal cellular behavior (**Supplementary Figure S2E-K**). DMSO had no significant impact on phagocytosis under confinement, but did slightly raise phagocytic cell number and decrease phagosome number per cell in media (**Supplementary Figure 2E-H**). With respect to motility, DMSO did not significantly affect velocity, distance traveled, or persistence under confinement or in media (**Supplementary Figure 2I-K**). We concluded based on these findings that DMSO is an appropriate vehicle control for our inhibitor treatment studies. Any significant decrease between DMSO vehicle and drug-treated cells under confinement can be accepted with reasonable confidence to be an effect of the inhibitor.

Arp2/3 complex disruption with the small molecule inhibitor CK-666 decreased the percent of phagocytic cells in both media and confined conditions compared to vehicle (**Figure 3A, quantified in Figure 3B).** Arp2/3 disruption decreased phagosome area under confinement (**Figure 3C, Supplementary Figure 2L**) and phagosome number in confined and unconfined settings (**Figure 3C, Supplementary Figure S2M**). Arp2/3 complex disruption also impaired phagocytic priming under confinement, as evidenced by decreased migratory persistence (**Figure 3D, Supplemental Movie 2**). Conversely, Arp2/3 complex disruption only decreased cell velocity and distance traveled in the media condition, but not migratory persistence after phagocytosis (**Supplementary Figure S2N-O**). We interpret these results to mean that Arp2/3 complex is crucial for phagocytic uptake, trafficking of targets to the phagolysosome, and phagocytic priming in confined settings. While phagocytic uptake was affected in the media condition, Arp2/3 loss of function in this context had less impact than it did under confinement. These results suggest that there may be varying degrees of compensation for loss of Arp2/3 complex function that are context-dependent.

**Figure 3:**
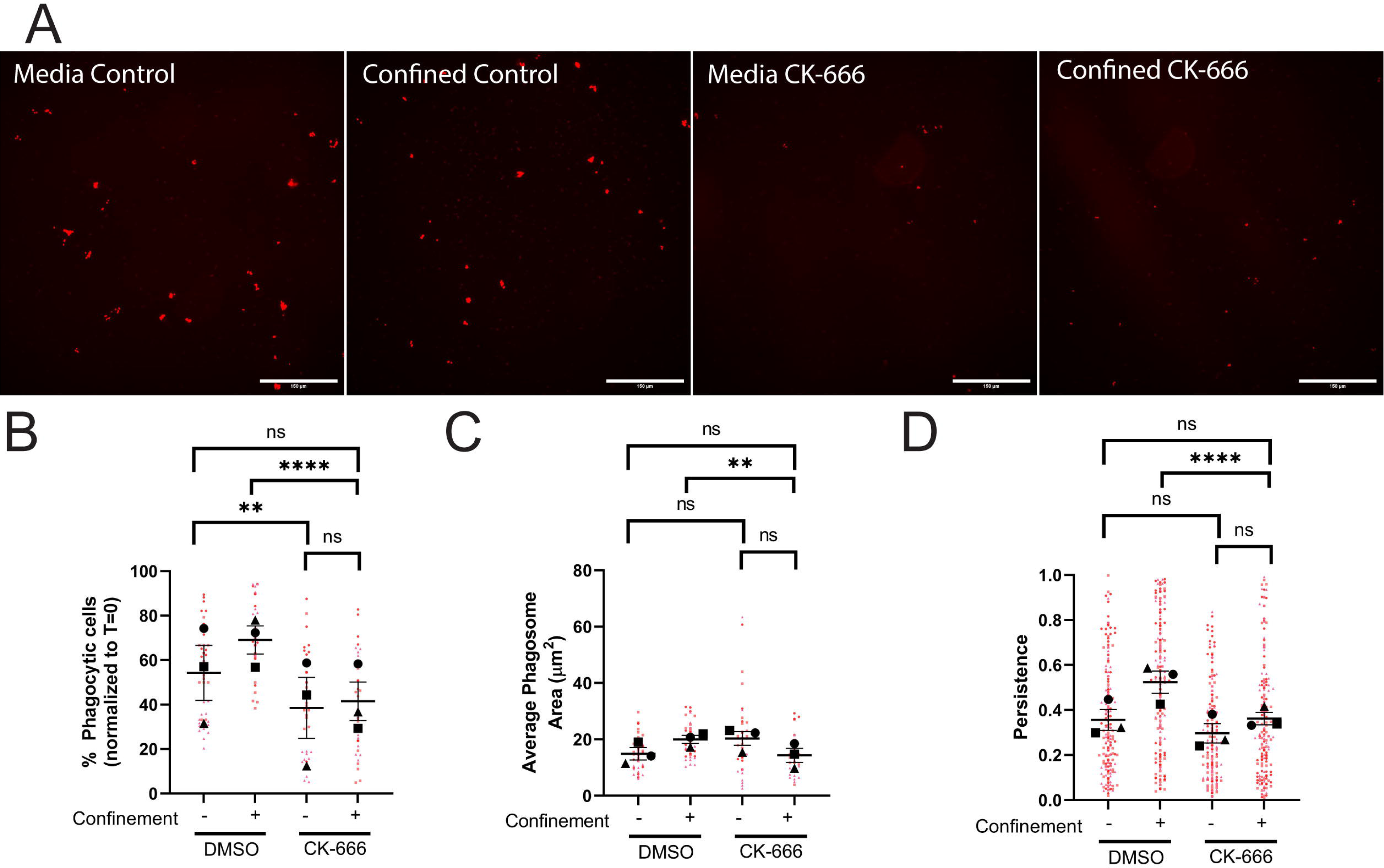
The Arp2/3 complex is required for phagocytosis and phagocytic priming in response to confinement. A-D) These cells were either treated with DMSO or 125µM CK-666. A) Example images of pHrodo-red label demonstrating internalized pHrodo-IgG-beads in each condition: Media control (vehicle – DMSO), Confined control (1% Agarose) plus Vehicle, Media control plus CK-666, Confined (1% Agarose) plus CK-666. Scale bar represents 100µm. B) The percentage of fluorescent cells in a field of view, normalized to T=0. C) Average phagosome size (µm^2^). D) The persistence of the cell during the length of its migration track. For all graphs, black points demonstrate experiment means, colored points demonstrate individual cell values for each run. N = 3 experiments for each graph; n = 15 fields of view for each condition per experiment for phagocytosis (B, C) or n = 50 cells for each condition per experiment for migration (D). Statistical analysis was assessed using Kruskal–Wallis with Dunn multiple comparisons: ns = not significant, **p < 0.01, ****p < 0.0001. Error Bars represent SEM.

### The actin cytoskeleton and non-muscle myosin II facilitate phagocytic priming and uptake

Previous studies examining myosin II in confined environments have demonstrated that it is needed for cell velocity but not for directional responses to either chemotactic (*30*) or haptotactic cues (*26*). We hypothesized in light of this that myosin II-dependent contractility would be involved in phagocytic uptake, but not phagocytic priming under confinement. Inhibition of myosin II with blebbistatin decreased phagocytic cell number in both confined and unconfined settings (**Figure 4A, quantified in Figure 4B**). However, confinement partially rescued phagocytosis in blebbistatin-treated cells, as judged by increased uptake in these cells compared to blebbistatin-treated cells in media. This points to myosin-independent cytoskeletal elements that are activated by confinement and compensate for loss of actomyosin contractility. Bead trafficking and phagosome formation in cells that take up beads were unaffected by blebbistatin treatment, as demonstrated by all phagosome measurements (**Figure 4C, Supplemental Figure S3A**). We interpret this to mean that myosin II impairment specifically affects phagocytic cup formation rather than intracellular trafficking to the phagolysosome.

**Figure 4:**
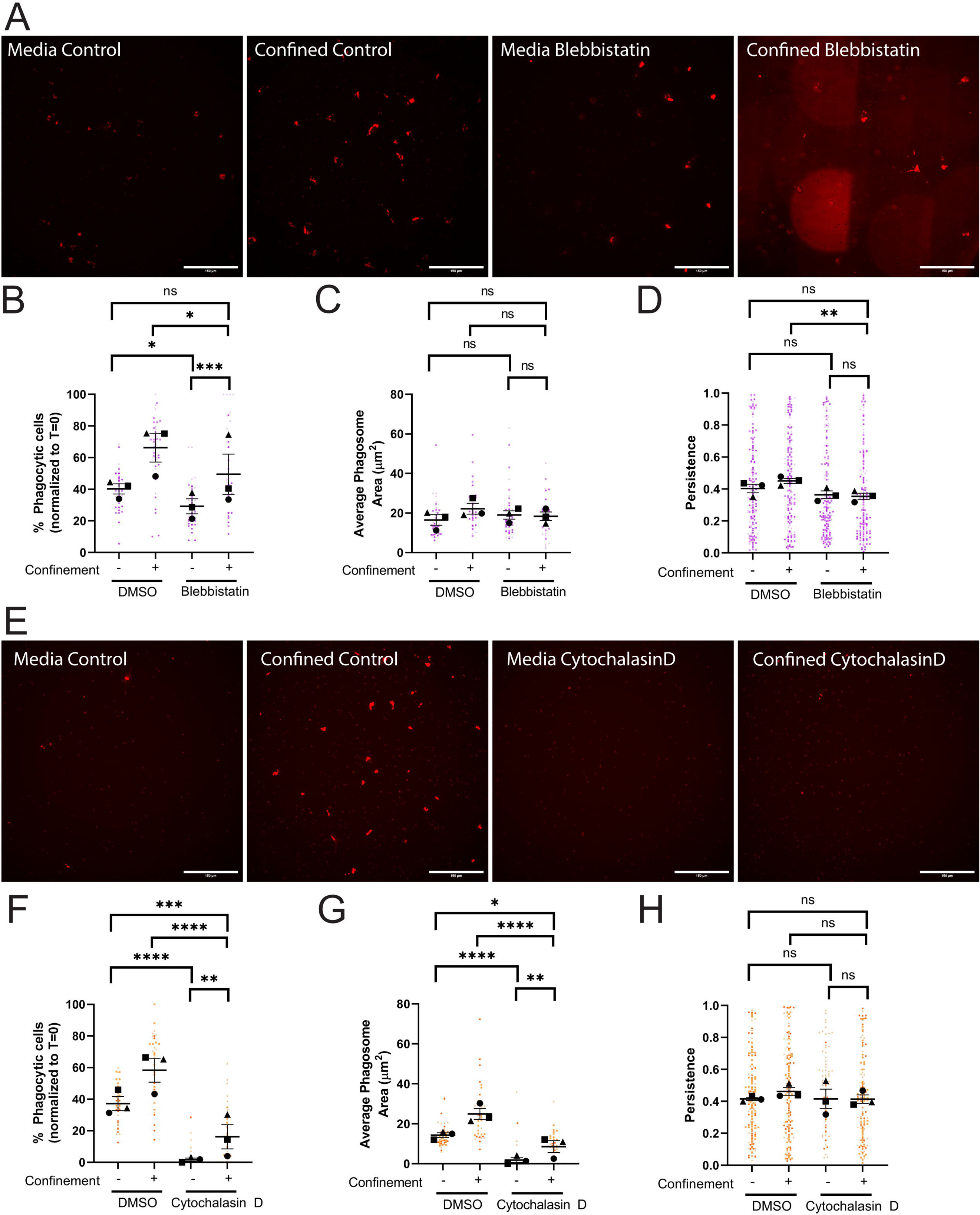
Confinement enhances phagocytic uptake and priming via actomysosin. A-D) These cells were either treated with DMSO or 30µM Blebbistatin. Blebbistatin caused strong auto-fluorescence when polymerized into agarose gels. These fluorescent fragments are visually distinct from beads in phase contrast. Composite images with both phase contrast and DsRed channels present were used to count phagocytic cells and manually remove blebbistatin autofluorescence from phagosome calculations. A) Example images of pHrodo-red label indicating internalized pHrodo-IgG-beads in each condition: Media control (vehicle – DMSO), Confined control (1% Agarose) plus Vehicle, Media control plus Blebbistatin, Confined (1% Agarose) plus Blebbistatin. Scale bar represents 100µm. B) The percentage of fluorescent cells in a field of view, normalized to T=0. C) Average phagosome size (µm^2^). D) The persistence of the cell during the length of its track. E-H) These cells were either treated with DMSO or 1µM Cytochalasin D. E) Example images of pHrodo-red label indicating internalized pHrodo-IgG-beads in each condition: Media control (vehicle – DMSO), Confined control (1% Agarose) plus Vehicle, Media control plus Cytochalasin D, Confined (1% Agarose) plus Cytochalasin D. Scale bar represents 100µm. F) The percentage of fluorescent cells in a field of view, normalized to T=0. G) Average phagosome size (µm^2^). H) The persistence of the cell during the length of its track. For all graphs, black points demonstrate experiment means, colored points demonstrate individual cell values for each run. N = 3 experiments for each graph; n = 15 fields of view for each condition per experiment for phagocytosis (B, C, F, G) or n = 50 cells for each condition per experiment for migration (D, H). Statistical analysis was assessed using the Kruskal–Wallis with Dunn multiple comparisons: ns = not significant, *p < 0.05, **p < 0.01, ***p < 0.001, ****p < 0.0001. Error Bars represent SEM.

Contrary to our expectations, myosin II disruption decreased phagocytic priming under confinement, as evidenced by decreased persistence of confined cells treated with blebbistatin (**Figure 4D, Supplemental Movie 3)**, while also decreasing velocity and distance traveled in this setting (**Supplemental Figure S3B**). Conversely, unconfined cells treated with blebbistatin showed no defect in post-phagocytosis persistence, again suggesting that myosin II’s contribution to phagocytic responses is context-dependent (**Figure 4D**). To investigate the impact of beads on the blebbistatin phenotypes, we repeated these experiments without beads. Blebbistatin-treated cells moved similarly to their DMSO treated counterparts in terms of velocity, persistence, and distance traveled when comparing within the confined and unconfined groups (**Supplementary Figure S3C**). These findings differ from our findings when pHrodo-IgG beads were present. Phagocytic uptake could cause cells to patrol their environments more deliberately, especially under confinement, in a much more myosin II-centric fashion.

Inhibition of either the Arp2/3 complex or myosin II did not fully impair phagocytosis under confinement, pointing towards another component playing a compensatory role during confined phagocytosis. We next treated cells with cytochalasin D to inhibit new actin filament formation, examining the necessity of filamentous actin (F-actin) in phagocytic uptake and priming under confinement. Cytochalasin D treatment resulted in a significant decrease in confined and unconfined phagocytosis (**Figure 4E**, **quantified in Figure 4F)**. Remarkably, confinement partially rescued phagocytosis in the presence of cytochalasin D. Phagosome formation, area, and number per phagocytic cell were all similarly affected by cytochalasin D, again with partial rescue by confinement (**Figure 4G, Supplementary Figure S3D**). Given the fundamental importance of actin in phagocytic cup formation and endocytic trafficking, these results are not unexpected. However, these data suggest that in confined conditions a less efficient actin-independent phagocytosis pathway may exist.

It was less straightforward to interrogate phagocytic priming during cytochalasin D treatment. Cytochalasin D-treated cells were dramatically less migratory, and traveled far lower distances than DMSO-treated counterparts in unconfined and confined settings (**Supplementary Figure S3E**). As already mentioned, cytochalasin D-treated cells were far less phagocytic than DMSO-treated counterparts (**Figure 4F**). Given these findings, it is difficult to interpret the lack of significance between DMSO- and cytochalasin D-treated persistence under confinement (**Figure 4H**). However, since myosin II and Arp2/3 complex inhibition affected phagocytic priming, it is reasonable to suspect that phagocytic priming would be impacted if F-actin assembly is dramatically compromised. Given that Arp2/3 complex (*31*) and myosin II (*32*) are both effectors of cell adhesion, we next interrogated the role of ECM-integrin engagement as an additional factor regulating phagocytic uptake and priming under confinement.

### Phagocytic priming, but not uptake, is dependent on ECM interaction

We started by using cilengitide to inhibit αv-containing integrins, which would be expected to impair BV-2 interaction with the fibronectin substrate on the glass surface in confined and unconfined settings. Cilengitide treatment did not affect phagocytosis in either condition, compared to vehicle control (**Figure 5A-B**). Overall, phagosome measures were similar between DMSO- and cilengitide-treated cells under confinement (**Figure 5C, Supplementary Figure S3F**). Thus, inhibition of a subset of αv-containing integrins did not impact uptake or intracellular trafficking of pHrodo-IgG-beads.

**Figure 5:**
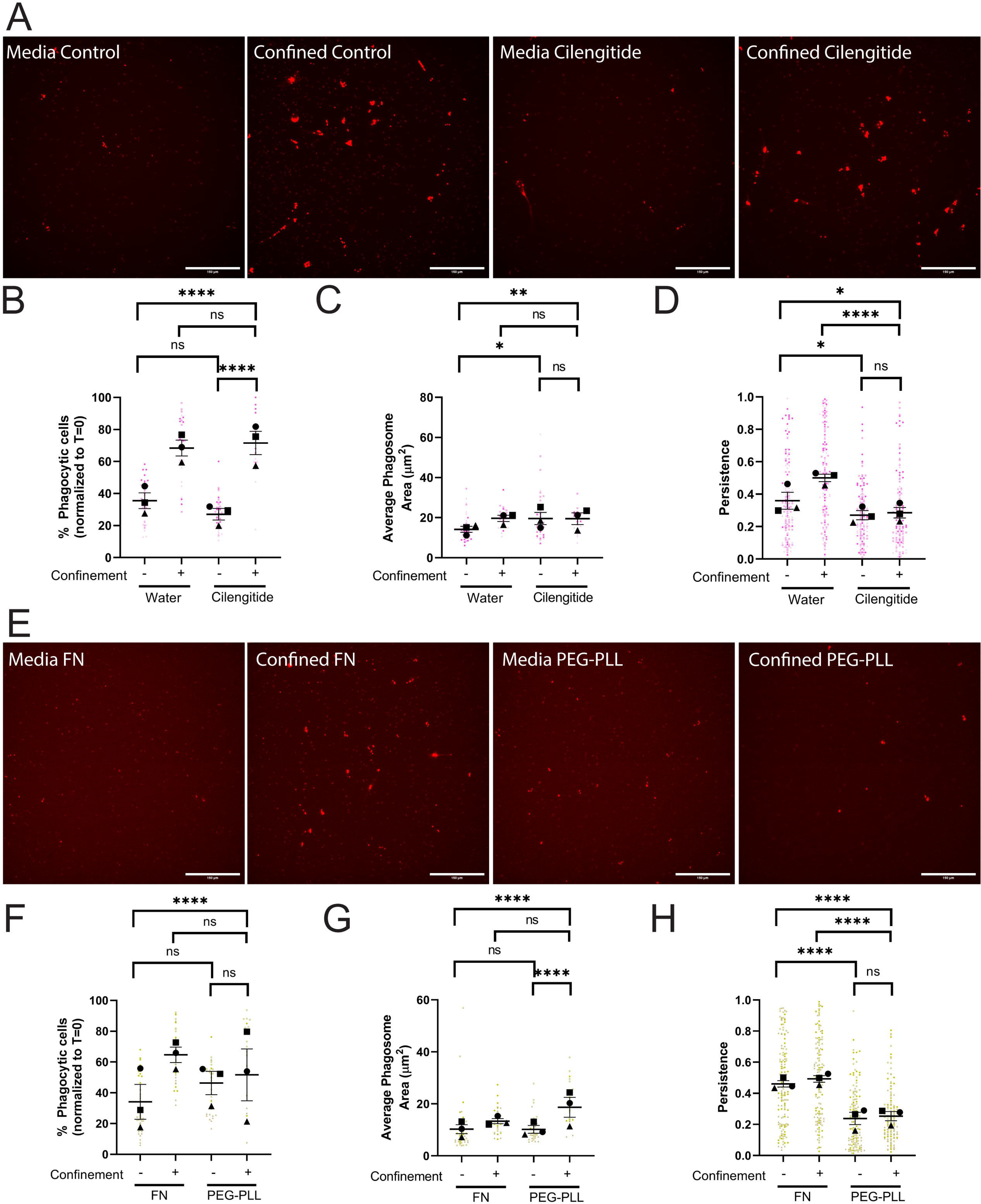
Cell adhesion is required for confinement induced phagocytic priming, but not uptake. A-D) These cells were either treated with water or 60µM Cilengitide. A) Example images of pHrodo-red label indicating internalized pHrodo-IgG-beads in each condition: Media control (vehicle – water), Confined control (1% Agarose) plus Vehicle, Media control plus Cilengitide, Confined (1% Agarose) plus Cilengitide. Scale bar represents 100µm. B) The percentage of fluorescent cells in a field of view, normalized to T=0. C) Average phagosome size (µm^2^). D) The persistence of the cell during the length of its track. E-H) These cells were either coated on 10µg/mL FN or 100µg/mL PEG-PLL. E) Example images of pHrodo-red label indicating internalized pHrodo-IgG-beads in each condition: Media control (vehicle – FN coating), Confined control (1% Agarose) plus Vehicle, Media control plus PEG-PLL coating, Confined (1% Agarose) plus PEG-PLL coating. Scale bar represents 100µm. F) The percentage of fluorescent cells in a field of view, normalized to T=0. G) Average phagosome size (µm^2^). H) The persistence of the cell during the length of its track. For all graphs, black points demonstrate experiment means, colored points demonstrate individual cell values for each run. N = 3 experiments for each graph; n = 15 fields of view for each condition per experiment for phagocytosis (B, C, F, G) or n = 50 cells for each condition per experiment for migration (D, H). Statistical analysis was assessed using the Kruskal–Wallis with Dunn multiple comparisons test correction: ns = not significant, *p < 0.05, **p < 0.01, ****p < 0.0001. Error Bars represent SEM.

However, cilengitide dramatically impacted phagocytic priming. Cilengitide-treated cells were less persistent than control cells in both settings (**Figure 5D, Supplemental Movie 5**). Our data also supports the idea that cilengitide-treated cells may be less adhesive, as treated cells migrate faster and move farther than control in both settings we assessed (**Supplementary Figure 3G**). These data suggest that phagocytic uptake and priming are not governed by identical pathways, as cilengitide-treated cells specifically have a defect related to post-phagocytosis migration rather than overall phagocytic ability. Since some integrin function is likely retained even in the presence of cilengitide, we were curious to see whether we could replicate these results by removing the response to ECM completely. To do this, we coated our dishes with 100 µg/mL PEG-Poly-L-Lysine (PEG-PLL) to limit integrin-mediated adhesion and compared cell responses to our standard coating of 10 µg/mL fibronectin. PEG-PLL elicits no change in phagocytosis compared to FN in either context (**Figure 5E-F, Supplementary Figure 3H**). However, phagocytic priming was dramatically impacted by PEG-PLL. In both contexts, post-phagocytic persistence was significantly impaired when cells were plated on PEG-PLL compared to FN (**Figure 5H**). Velocity and distance traveled increased on PEG-PLL under confinement, but not in media (**Supplementary Figure S3I**). All of these results were consistent with the results of cilengitide treatment, and demonstrated that ECM engagement was a key factor in phagocytic priming, especially under confinement, but not in phagocytic uptake.

## DISCUSSION

In the present study, we report that confinement increases phagocytosis, enabling more cells to take up IgG-opsonized beads compared to their unconfined counterparts. In addition, we discovered that phagocytosis had a “priming” effect on cell motility, resulting in increased cell persistence after cells underwent an initial phagocytic event. Phagocytic priming was especially pronounced under confinement. Confined phagocytosis resembled unconfined phagocytosis, except that confinement is capable of partially rescuing phenotypes seen in unconfined settings. This led us to propose a two-part process that allows phagocytes to respond to phagocytic cues under confinement. First, confinement stimulates Arp2/3 complex and myosin II to take up IgG-beads more efficiently (**Figure 6**). Then, particle internalization primes phagocytes to migrate persistently, which requires Arp2/3 complex, myosin II and ECM engagement (**Figure 6**). Our findings suggest that coherent interplay between mechanical sensing, ECM sensing, and phagocytic uptake determines cellular sensing and response to phagocytic targets. Some of these factors influence specific elements of this process, and the interplay between them and the cytoskeleton is worth exploring further.

**Figure 6:**
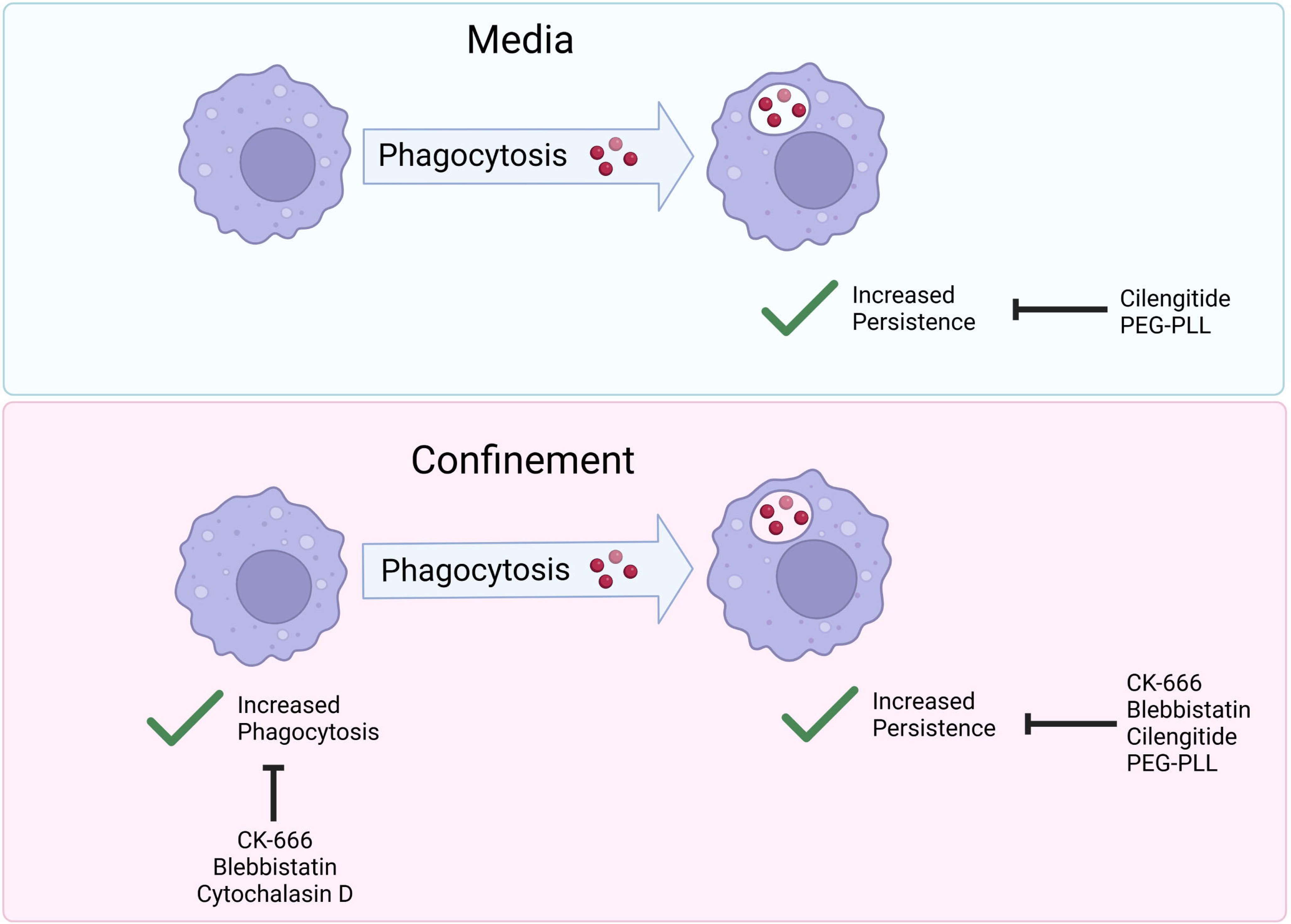
The actin cytoskeleton and cell-ECM adhesion respond to confinement and phagocytic targets to tune phagocytic uptake and priming. Graphical Abstract summarizing paper findings. Cells in media experience increased persistent migratory patterns. These patterns are disrupted when integrin binding is blocked (by Cilengitide), or fibronectin is replaced by PEG-poly-L-lysine (PEG-PLL). Cells under confinement experience increased phagocytosis and increased phagocytic priming. Increased persistence is mediated by myosin II (inhibited by blebbistatin), Arp2/3 complex (inhibited by CK-666) and integrin-ECM engagement (inhibited by cilengitide or PEG-PLL). Increased phagocytosis is mediated by Arp2/3 complex and myosin II (inhibited by CK-666 and blebbistatin, respectively).

Post-phagocytic cells under confinement consistently demonstrated directionally persistent migration, which we term phagocytic priming. Disrupted phagocytic priming sometimes correlated with altered phagosome measurements (treatment with CK-666 or cytochalasin D), but other times phagosome measurements were unaffected by the lack of priming (treatment with blebbistatin, cilengitide, or cells plated on PEG-PLL). A study by Procès *et al* found that BV-2 cells activated via mechanical stretch became more directionally persistent in one-dimensional assays in PDMS microchannel devices (*33*), suggesting that mechanotransduction of external forces by the actomyosin cytoskeleton may be the underlying driver of phagocytic priming. This is in line with the results of this study, as key drivers of phagocytic priming identified in this study, myosin II and The Arp2/3 complex, are responsive to mechanical force (*30*, *34–38*). In addition, stretch-induced persistent migration of BV-2 cells activated Arp2/3 complex in lamellipodial protrusions via Rac GTPases (*33*). Procès *et al* also noted increased persistence of BV-2 cells after treatment with lipopolysaccharides (LPS) compared to mechanically activated counterparts, with both activation methods being more persistent than control cells (*33*). Previous studies have also demonstrated increased migratory persistence under confinement independent of an external gradient when examining cell migration in 3D environments using cancer cells (*39*, *40*) or fibroblasts as model systems (*41*). These findings align well with our data, suggesting that persistent migration is inducible by physical force, and that pathogen sensing similarly cooperates with mechanical confinement to alter motility.

IgG opsonizes foreign bodies in the context of infection, and can initiate tissue inflammatory immune pathways by binding to IgG through the Fc gamma receptors (FcγRs) expressed on the plasma membrane (*42*). Phagocytosis of IgG-bound particles aids in clearance of pathogens and foreign debris. One example of this *in vivo* is immature dendritic cells that surveil tissue until they encounter an antigen (possibly bound by IgG), at which point they mature and follow a gradient of CCL21 to the lymph nodes in order to incite a T cell response to the pathogen (*43*, *44*). Phagocytic priming could enable cells to move more directionally to the lymphatics, as well as better managing pathogen control in peripheral tissues. The impact of IgG on microglial function in the central nervous system is complicated. Un-complexed IgG is proposed to have neuroprotective effects, via stimulation of microglial endocytosis (*45*). However, IgG-ligand complexes (as modeled by our IgG beads) have been proposed to exacerbate detrimental microglial responses in multiple sclerosis (*46*), while IgG-amyloid beta complexes are commonly present within amyloid plaques (*47*). It is possible that Arp2/3-dependent, ECM-responsive haptotaxis helps explain the response to these IgG complexes, as it has been implicated previously in haptosensing (*28*). Phagocytic priming may help explain how microglia find these IgG complexes, and why they remain associated with lesions and plaques *in situ*. Future studies will also reveal whether phagocytic priming is a universal response to phagocytosis or unique to IgG. It is also imperative to examine whether primed cells are more responsive to chemotactic or haptotactic cues.

Paterson and Lämmermann recently discovered that macrophages manage phagocytosis via an integrin-dependent process termed haptokinesis (*25*). When following cellular behavior in a 3D Matrigel context, removal of β1 integrin or talin significantly impaired motility, which disrupted macrophages’ ability to come into contact with phosphatidylserine beads or apoptotic cells to internalize them. Of note, they also demonstrated that β1 integrin or talin disruption did not impair phagocytosis when cells were in suspension. Consistently, we demonstrate no defect in phagocytic uptake upon cilengitide treatment or removal of fibronectin. Furthermore, we also show that there is an impact on motility in the context of phagocytic priming. These results are largely consistent with the previous study, with the caveat that cells in our studies have more freedom to interact with beads compared to 3D Matrigel. In addition, our motility measurements were focused on post-phagocytic behavior, which necessarily selects for cells that have successfully taken up beads regardless of treatment.

The phagocytic priming response suggests that cells may be re-orienting their microtubule organizing center (MTOC) after bead internalization. MTOC re-orientation correlates with polarized motility, which makes this an interesting future direction for examining the intersection of confinement and phagocytic priming on cells (*48*). MTOC disruption leads to decreased cell polarity and cell migration, leading to less persistent cancer cell motility (*49*). Similar findings have been demonstrated for fibroblasts (*50*) and mouse melanoma cells (*51*) as well. Another aspect relevant to the present study is the finding that microtubules are important for mechanosensing in astrocytes (*52*). Increasing substrate rigidity resulted in polarized microtubule creation, which in turn enhanced focal adhesion function to promote actomyosin contractility (*52*). At the most fundamental level, microtubules are required for both phagocytic uptake and phagosome maturation (*53*). These results together suggest that MTOC and microtubule polarization could fundamentally link phagocytic uptake to polarized post-phagocytosis migration. In addition, mechanosensitive microtubule signaling might be a factor that contributes to confinement partially rescuing phagocytosis upon cytochalasin D treatment.

The present work is consistent with the idea that integrating microenvironmental factors in a controllable fashion is key to understanding complex cellular behaviors. The current work reveals an intersection of mechanical cues, adhesion cues, and phagocytic uptake together stimulate a two-part response to IgG beads, driven by differential cytoskeleton regulation. This integration of multiple microenvironmental factors is important in aligning a cell’s behavior in an *in vitro* setting with its *in vivo* function. Over time it will be possible to model microglia *in vitro* in a way that more completely captures their *in vivo* morphology and behavior, which is important to further understand their role in neurological diseases and to reveal future therapeutic targets (*45–47*, *54–58*).

## MATERIALS AND METHODS

### Cells

The murine microglia-like BV-2 cell line was used for all experiments. Cells were grown in complete cell culture media, containing Dulbecco’s Modified Eagle Medium (DMEM) (Gibco, 31053028), 5% fetal bovine serum (FBS) (Gibco, 10437028), 1% glutaMAX (Gibco, 35050061), and 1x Antibacterial-Antimycotic (Gibco, 15240062) at 37°C and 5% CO_2_. 70-80% confluent dishes were treated with 0.05% Trypsin-EDTA solution (Caisson Labs TRL02-100ML) at 37°C for 10 minutes. The solution was then aspirated and replaced with 2mL of complete cell culture media that was gently sprayed over the bottom of the dish to dislodge cells and collect for counting and passage into new dishes.

### Four well chamber preparation

4 well chambers (Cellvis, C4-1.5H-N) were coated in 10μg/mL fibronectin (Gibco, 33016015) for 1 hour at 37°C. Chambers were washed three times with 1x Cell Culture Phosphate Buffered Saline (Corning, MT21040CV) (referred to from here on as PBS). Two wells were each filled with 1mL PBS and two wells were each filled with 1mL 1% agarose. <u>Agarose preparation</u>: 2.5mL of 2x Hank’s Balanced Salt Solution (Sigma, H1387-10X1L) and 5mL of complete cell culture media (as defined above) were combined and placed in a 68°C bead bath for 1 hour. 0.12g of low gelling temperature agarose (Sigma, A9045-5G) was added to 2.5mL of sterile water (Corning, 25-055-CV) and heated in a microwave until fully dissolved. The two solutions were combined to create a warm 1% agarose mixture. 1mL of the mixture was poured into each well and allowed to solidify for 90 minutes at room temperature. Chambers were wrapped in parafilm and stored at 4°C for up to one month.

### Phagocytosis bead preparation

100µg of Normal Mouse IgG (Sigma-Aldrich, 12-371) was pHrodo labeled using the pHrodo™ iFL Red Microscale Protein Labeling Kit (ThermoFisher, P36014) according to manufacturer instructions. 60µL of pHrodo-IgG was opsonized to 60µL of 2-micron Polybead Carboxylate Microspheres (Polysciences, Inc, 18327-10) (hereafter referred to as beads) in 3mL of 1x PBS at 37°C for one hour. Beads received three washes with 1x PBS after completing opsonization and were resuspended in a final volume of 500µL, stored at 4°C for up to 4 months. Before usage in an experiment, beads were vortexed vigorously.

### Cell preparation

Agarose wells had two punches placed in each for cell and bead insertion using a 0.75mm biopsy punch tool (LabTech, 52-004908). Confluent dishes were treated according to the standard passaging protocol (above). For cell migration experiments, 13,000 cells were added to each media well. For phagocytosis, 18,000 cells were added to each media well. Lower numbers were used for migration experiments to minimize cell-cell interactions and allow more room for uninterrupted random migration. For both types of experiments in the confined condition, 15,000 cells were inserted under the agarose in each punched location using a gel-loading pipette tip. If the amount of media containing 15,000 cells exceeded 15µL, the cells were spun down at 1000x g and resuspended in 10µL of complete cell culture media. Chambered dishes were placed at 37°C for three hours to allow for cells to sit down and spread before being moved to the microscope. For the inhibitor trials, each cell culture media well received either vehicle or inhibitor to create the working concentration used for cell treatment (more information below).

### Phagocytosis microscope assay

Upon completing the 3-hour incubation, IgG beads were added to wells. Media wells received 5µL of beads. For confined wells, a 1:4 dilution of the beads was made with PBS and inserted into each punch in the agarose to match bead density with the media condition. Chambers were then placed in a Tokai Hit INU incubation system controller on the Olympus IX83 and maintained at 37°C, 90% humidity, and 5% CO_2_ for the duration of the live cell imaging. A 20x air objective was employed during overnight time lapse imaging. 15 fields of view per well were selected. Images were taken at each field of view every 10 minutes for the span of 8 hours. Both the relief contrast and the DsRed channels were utilized. The DsRed channel was set to 500 ms exposure to detect pHrodo signal upon bead internalization by cells.

### Analyzing bead density

Each phagocytosis file was opened in Fiji ImageJ. A bandpass filter was then run on the relief contrast channel. Structures were filtered to fit between 5 and 15 pixels and then threshold filtered until only the beads were highlighted in red. The analyze particles function was used to count bead density. Area threshold was set to 10-30 micron^2^, circularity threshold was set to 0.8-1.0, and outlines were shown. When summarized, average size was maintained between 15 and 17 micron^2^ across the time lapse, with threshold being adjusted and analysis rerun if the averages were too small or too large. Counts of beads per time point were averaged to create the bead density designator for each file. Averaged bead numbers falling between 50 and 500 were designated as low bead density; between 500 and 950 as medium bead density; and between 950 and 1350 as high bead density.

### Analyzing phagocytic rate

Each phagocytosis file was opened in FIJI ImageJ. Channel colors were corrected and brightness/contrast adjusted to create the same brightness and contrast parameters for each file. Using the cell counter plugin, the total number of fluorescent and non-fluorescent cells was counted at each measured time point throughout the phagocytosis videos. Comparisons between treatment groups at individual time points as well as comparing changes over time within each group were conducted with GraphPad Prism. Graphs were normalized to T=0. Fluorescent cells at the beginning of the videos were removed from the count of both fluorescent cells and total cells per field of view. Data comparisons were broken up by bead density, only comparing media and confined fields in the low bead density category to each other. Blebbistatin auto-fluoresces under fluorescent imaging. Visual inspection of phagocytic cells ensured that counted fluorescent internalized pHrodo-IgG and not blebbistatin.

### Analyzing phagosome size

Each phagocytosis file was opened in CellSens Dimension software under the Count and Measure plugin. Under detection options, minimum object size was set to 5 pixels. Under the Thresholding tab, the adaptive threshold was selected. Selecting the DsRed channel, the maximum was set to 65000 (the maximum) and the minimum was changed for each image to label only internalized beads, although never going below 1000. Selecting the relief contrast channel, the maximum was set to 2 and the minimum to 1 so that only fluorescent labels were detected. Outputs of total number of phagosomes and their average size per field of view were then verified at each timepoint before the data was exported to an excel file. This data was presented three ways. The average size of phagosome was reported by dividing the average size per field of view at 2 hours by the total number of phagosomes detected at 2 hours. Average size of phagosome per phagocytic cells was reported by dividing the average size per field of view by the number of phagocytic cells at that timepoint. The average number of phagosomes per phagocytic cell was reported by dividing the total number of phagosomes at 2 hours by the number of phagocytic cells at that timepoint. Average size of phagosome and average size of phagosome per phagocytic cell differ as the former examines whether beads are trafficked into the same or different lysosomes and the latter examines if more beads overall are being trafficked in each treatment condition.

### Analyzing motility

Each motility file was opened in FIJI ImageJ. The Manual Tracking plugin was used to track 10 cells per field of view and combining data across all fields of view within treatment groups. Cell tracking was stopped if a cell divided or ran into another cell, resulting in a changed direction. The resulting measurements were uploaded into the Ibidi Chemotaxis and Migration Tool ImageJ plugin to measure velocity, accumulated distance, and persistence. Persistence (also termed D/t) is a measure of the straight-line distance between a cell’s starting and ending position on a track (D) divided by its overall track length (t). Thus, a value very close to zero (high t, low D) is understood to be meandering much more than a cell moving along the shortest distance between its start and end (d = T), which has a value of 1. The closer the value to 1, the more persistent the migration is. These data tables were exported to GraphPad Prism to graph differences in velocity, distance, and persistence between treatment groups.

### Different agarose concentrations

4 well chamber dishes were coated with fibronectin as described above. These chambered dishes were made to contain one well of PBS, one well of 1% agarose, one well of 2.5% agarose, and one well of 5% agarose. 1% agarose was prepared as stated above. For both the 2.5% and the 5% agarose concentrations, only the amount of agarose added to the sterile water differed in the protocol. 2.5% agarose was prepared using 0.3g of agarose in 2.5mL of sterile water and 5% agarose was prepared using 0.6g of agarose in 2.5mL of sterile water. Chambers were wrapped in parafilm and stored at 4°C for up to one month.

### Microscope setup for motility assay

Upon completing the 3-hour incubation, chambers were placed in an environmental chamber on a Tokai Hit INU incubation system controller on the Olympus IX83 and maintained at 37°C, 90% humidity, and 5% CO_2_ for the duration of the live cell imaging. A 20X air objective was employed during overnight time lapse imaging. 10 fields of view per well were selected. Images were taken at each field of view in the relief contrast channel every 10 minutes for the span of 16 hours.

### Inhibitor-containing agarose

CK-666 (abcam, ab141231), Blebbistatin (Fisher Scientific, NC0664123), and CytochalasinD (ThermoFisher, PHZ1063) were resuspended in anhydrous DMSO (ThermoFisher, D12345) to create stock solutions. Cilengitide (Sigma Aldrich, SML1594-5MG) was resuspended in water (Corning, 25-055-CV) to create a stock solution. Agarose was prepared as previously described and then split into two aliquots of 5mL warm agarose solution. Each aliquot received either vehicle (anhydrous DMSO or water) or the drug stock solution at a predetermined concentration (CK-666 125μM, Blebbistatin 30μM, CytochalasinD 1μM, Cilengitide 60μM). Agarose chambers were prepared as previously described, wrapped in parafilm, and stored at 4°C for up to one month. For PEG-Poly-L-Lysine (PEG-PLL) (Creative PEG Works, PPL-2k20k3.5-100mg) plates, 2 wells were coated with 100μg/mL PEG-PLL for 1 hour at room temperature, then 10μg/mL fibronectin were added to the 2 non-coated wells and coated at 37°C for 1 hour. Agarose was then prepared as previously described.

### Inhibitor phagocytosis and motility runs

Cells were prepared as previously described. Media wells contained either vehicle or drug at the same concentration present in the agarose. Cells were seeded as previously described and plates run on Olympus IX83 as previously detailed.

### Statistical Analysis

The Kruskal–Wallis with Dunn multiple comparisons test was used to assess significance in experiments where a normal distribution of the dataset could not be assumed. When only two experimental conditions were tested, we used Mann–Whitney tests when we could not assume normality. Unpaired t tests and ANOVAs were used when normality tests indicated a normal distribution of the data. All statistics were calculated using GraphPad Prism, and significance was assumed if p ≤ 0.05. More information on each statistical test can be found in the relevant figure legend panel.

---

**Supplementary Figure 1:**
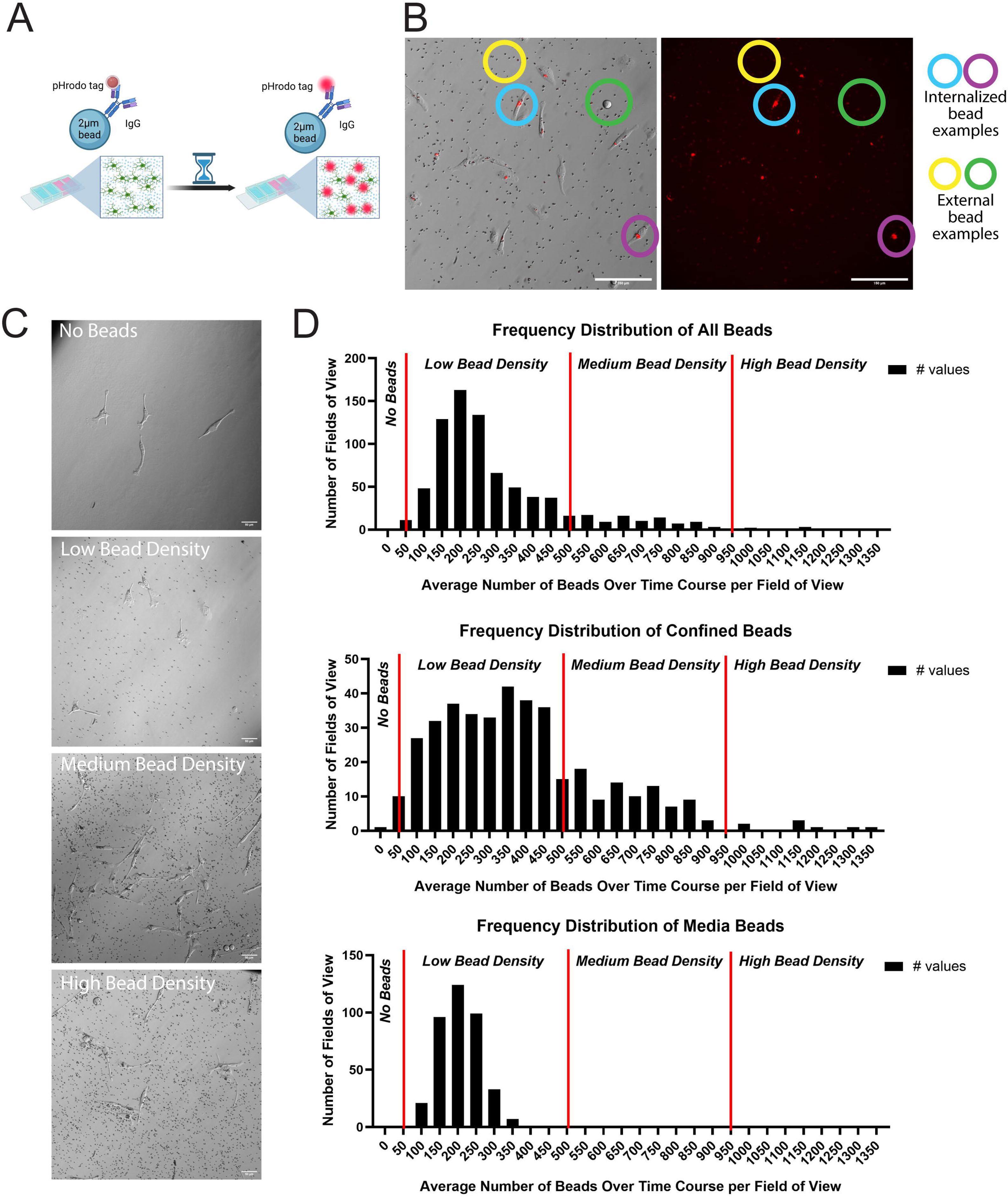
Strategy for controlling for bead density. A) Schematic depicting phagocytic bead labelling and the pHrodo tag fluorescing inside cells but not externally. B) Duplication of Figure 2A. Examples of either internalized (blue and purple circles) or external (yellow and green circles) pHrodo-red staining outlined in both composite and pHrodo only images. Scale bar represents 100µm. C) Example phase contrast images displaying different bead densities. No beads (top) through high bead density (bottom). Scale bar represents 50µm. D) Histograms detailing the breakdown of average bead densities per field of view across all experimental runs. Low bead density was classified as any field of view with a bead average between 50 and 500 beads; medium bead density was 500 to 950 beads; high bead density was any field of view average above 950 beads. Low bead densities in confined images most closely matched the density of typical media images.

**Supplementary Figure 2:**
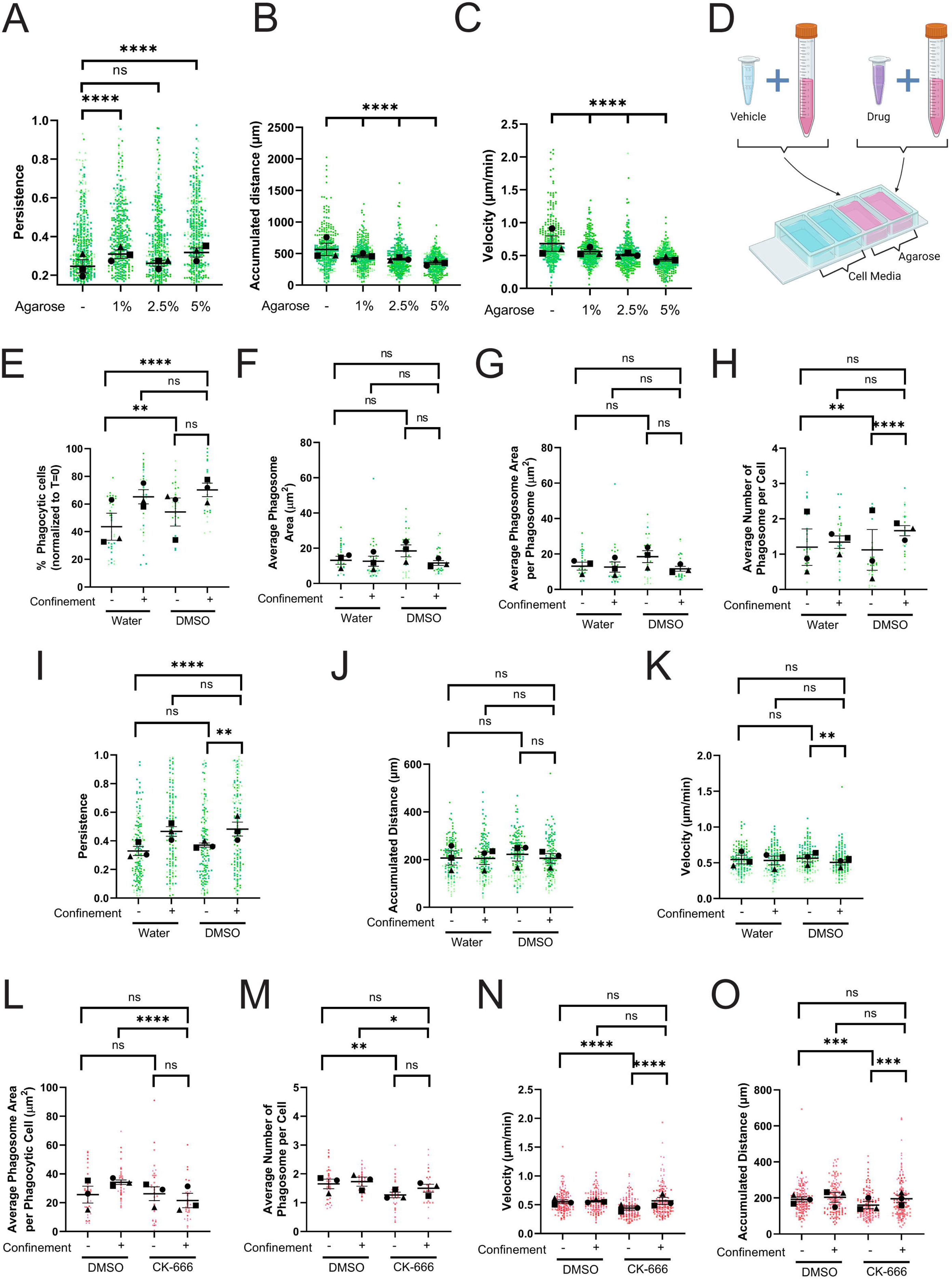
Controlling for agarose stiffness and DMSO influence on motility and phagocytosis. A) Persistence of the cells moving randomly (e.g. not post-phagocytic) in media, under 1% agarose, 2.5% agarose, or 5% agarose. B) Velocity (in µm/min) of cells moving in media, under 1% agarose, 2.5% agarose, or 5% agarose. C) Maximum total accumulated distance traveled (µm) for cells moving in media, under 1% agarose, 2.5% agarose, or 5% agarose. D) Schematic depicting how drugs were added into the agarose. Aliquots of 1% agarose were created and then vehicle or drug was added at the corresponding molarity before the agarose was poured into the confinement wells. E-K) These cells were either treated with water or DMSO. E) The percentage of fluorescent cells in a field of view, normalized to T=0. F) Average phagosome size (µm^2^). G) Average phagosome area per phagocytic cell (µm^2^). H) Average number of phagosomes per phagocytic cell. I) The persistence of the cell during the length of its track. J) Velocity of the cells migrating (µm/min). K) The maximum accumulated distance (µm) that the cell traveled during its tracking. L-O) These cells were either treated with DMSO or CK-666. These graphs relate to the experiments presented in Figure 3. L) Average phagosome area per phagocytic cell (µm^2^). M) Average number of phagosomes per phagocytic cell. N) Velocity of the cells migrating (µm/min). O) The maximum accumulated distance (µm) that the cell traveled during its tracking. For all graphs, black points demonstrate experiment means, colored points demonstrate individual cell values for each run. N = 3 experiments for each graph; n = 15 fields of view for each condition per experiment for phagocytosis (E-H, L-M) or n = 80 cells for each condition per experiment for migration (A-C, I-K, N-O). Statistical analysis was assessed using the Kruskal–Wallis with Dunn multiple comparisons test correction: ns = not significant, *p < 0.05, **p < 0.01, ***p < 0.001, ****p < 0.0001. Error Bars represent SEM.

**Supplementary Figure 3:**
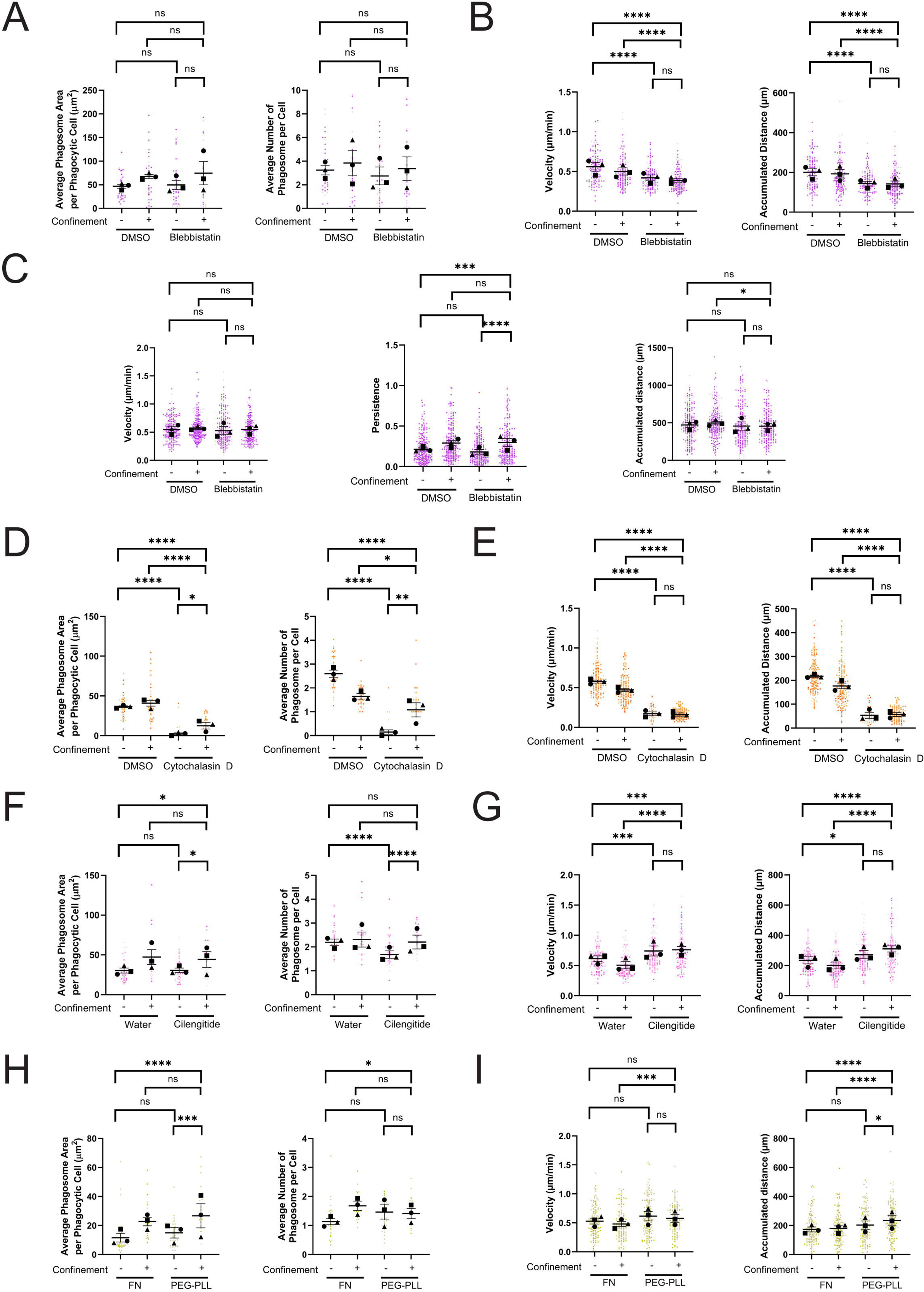
Additional phagocytic uptake and cell motility data related to Figures 4 and 5. A, B, D, E) These graphs relate to the experiments presented in Figure 4. F-I) These graphs relate to the experiments presented in Figure 5. A, D, F, H) Average phagosome area per phagocytic cell (left) and average phagosome number per phagocytic cell (right) when treated with the respective drug (Blebbistatin, Cytochalasin D, Cilengitide) or when plated on PEG-PLL. B, E, G, I) Velocity (µm/min) (left) and maximum accumulated distance traveled (µm) (right) of cells when treated with the respective drug (Blebbistatin, Cytochalasin D, Cilengitide) or when plated on PEG-PLL. C) Persistence (left), velocity (µm/min) (middle), and maximum total accumulated distance traveled (µm) (right) readings for cells with and without blebbistatin treatment when no beads are present. For all graphs, black points demonstrate experiment means, colored points demonstrate individual cell values for each run. N = 3 experiments for each graph; n = 15 fields of view for each condition per experiment for phagosomes (A, D, F, H) or n = 50 cells for each condition per experiment for migration (B, E, G, I). Statistical analysis was assessed using the Kruskal–Wallis with Dunn multiple comparisons test correction: ns = not significant, *p < 0.05, **p < 0.01, ***p < 0.001, ****p < 0.0001. Error Bars represent SEM.

## ACKNOWLEDGEMENTS

We thank the members of the Rotty Lab for helpful discussions during research development. This project is sponsored by the Uniformed Services University of the Health Sciences (USU); however, the information or content and conclusions do not necessarily represent the official position or policy of, nor should any official endorsement be inferred on the part of, USU, the Department of Defense, or the U.S. Government.

## AUTHOR CONTRIBUTIONS

S.P.: experiments and planning, data analysis, writing and editing. S.L.: experiments and planning, data analysis, editing. J.R.: Project oversight, experiments and planning, writing, editing, funding. All authors had the opportunity to review and comment on the manuscript prior to submission.

## DATA AVAILABILITY STATEMENT

All primary data will be openly available upon request.

